# Genedex: Contextualizing gene signatures across Alzheimer’s disease pathology, cell types, and brain regions

**DOI:** 10.64898/2026.10.26.761919

**Authors:** Giulia Pegoraro, Mehmet Enis Isgoren, Livia C Rizzo, Larisa Soto Morales, Pourya Naderi Yeganeh, Inyoung Sung, Sumeet Gupta, Doo Yeon Kim, Rudolph E Tanzi, Katie Lunnon, Isabel Castanho, Winston Hide

## Abstract

Systematic prioritization of candidate Alzheimer’s disease (AD) genes and their target functions for therapeutic intervention requires a consistent understanding of their brain location, cell-type-specific expression, relationships with pathology, and direction of change. Defining the role of a gene in AD requires interrogation of multiple publications, each providing a different context. Often, it is also unclear whether a gene is up-or downregulated across these contexts. However, the narrow diversity of contexts in which they have been assayed and described limits interpretation. We integrated 1,562 gene signatures from 34 studies, annotated for pathological process, cell type, brain region, and direction of change, and showed that Genedex recovers established AD biology. Genedex addresses cellular and regional unexplained directional contradictions from 10,657 discordant genes in the published AD literature. We investigated the role of heat-shock chaperones and neurotrophic signaling in cognitive resilience using Genedex. Heat-shock chaperone genes are broadly upregulated with tau burden; but, in the presence of amyloid pathology, their expression diverges, showing upregulation in excitatory neurons and downregulation in glia and during Aβ clearance. *LINGO1*, a regulator of BDNF, is upregulated with neuritic plaque load, tau burden, and selective vulnerability in cortical regions. Notably, in tau-affected *RORB*⁺ excitatory neurons, *LINGO1* expression is inversely correlated with neurotrophic genes *BDNF* and *NTRK2.* A prioritized interneuronal resilience signature shows correlated pathway co-expression with somatostatin and parvalbumin interneuronal signatures. Vulnerability-associated signatures separate by cellular context: neuronal programs are positively correlated with vulnerable interneurons, while glial and vascular cells are predominantly anti-correlated. By systematically mapping genes across functional, cellular, spatial, and directional axes, Genedex provides a platform for discovery of contextualized targets and mechanistic hypotheses. Genedex is freely available at hidelab.github.io/genedex-app.

## Introduction

A central question in Alzheimer’s disease (AD) research is whether a gene is associated with the disease and, if so, with which of its pathogenic processes. AD-associated processes can be described at different levels. Based on the neuropathological burden of amyloid-β (Aβ) plaques and tau tangles. AD can be classified into specific stages, such as using the Thal phases of Aβ deposition and Braak staging system (1,2). Based on cognitive status relative to pathology burden, the concept of cognitive resilience – the preservation of cognition despite neuropathology – has challenged neuropathology-focused definitions of AD (3). At the molecular level, changes in the brain include disease-associated glial states (4,5) and vulnerable or resilient neuronal programs (6). Together, these molecular, cellular, and neuropathological features form a continuum, representing the range of biological changes associated with AD progression (7).

Placing a gene in the context of these processes is challenging for three main reasons. First, AD is heterogeneous. For example, Aβ, tau, glial, and vascular processes interact over the decades before clinical onset (8,9). Second, gene signatures are usually annotated with general-purpose ontologies such as the Gene Ontology (10), which describe broad cellular processes rather than disease-specific mechanisms and need dedicated curation to capture AD mechanisms (11). Third, data-driven approaches are fragmented because published differentially expressed genes and co-expression modules (12) vary with cohort, brain region, case definition, and analysis (13,14). Defining the role of a gene in AD requires interrogation of multiple publications, each providing a different context. It is also often unclear whether a gene is up-or downregulated across these contexts.

Consistently placing a gene within the AD continuum requires integration of structured, disease-specific annotations with data-driven evidence, where each observation specifies the functional process (e.g., amyloid load), the cell type (e.g., microglia), the direction of change, and the tissue type (e.g., cortical tissue). All four dimensions are necessary because a gene may show opposite expression directions across cell types, regions, or processes. Existing resources capture only subsets of these dimensions. For example, AD-focused platforms such as AlzGPS (15), AlzData (16), Agora (17), and ssREAD (18) organize omics data by gene, cell type, or brain region. General-purpose resources use broad functional terms; however, none captures all four dimensions (Table 1).

**Table 1.**
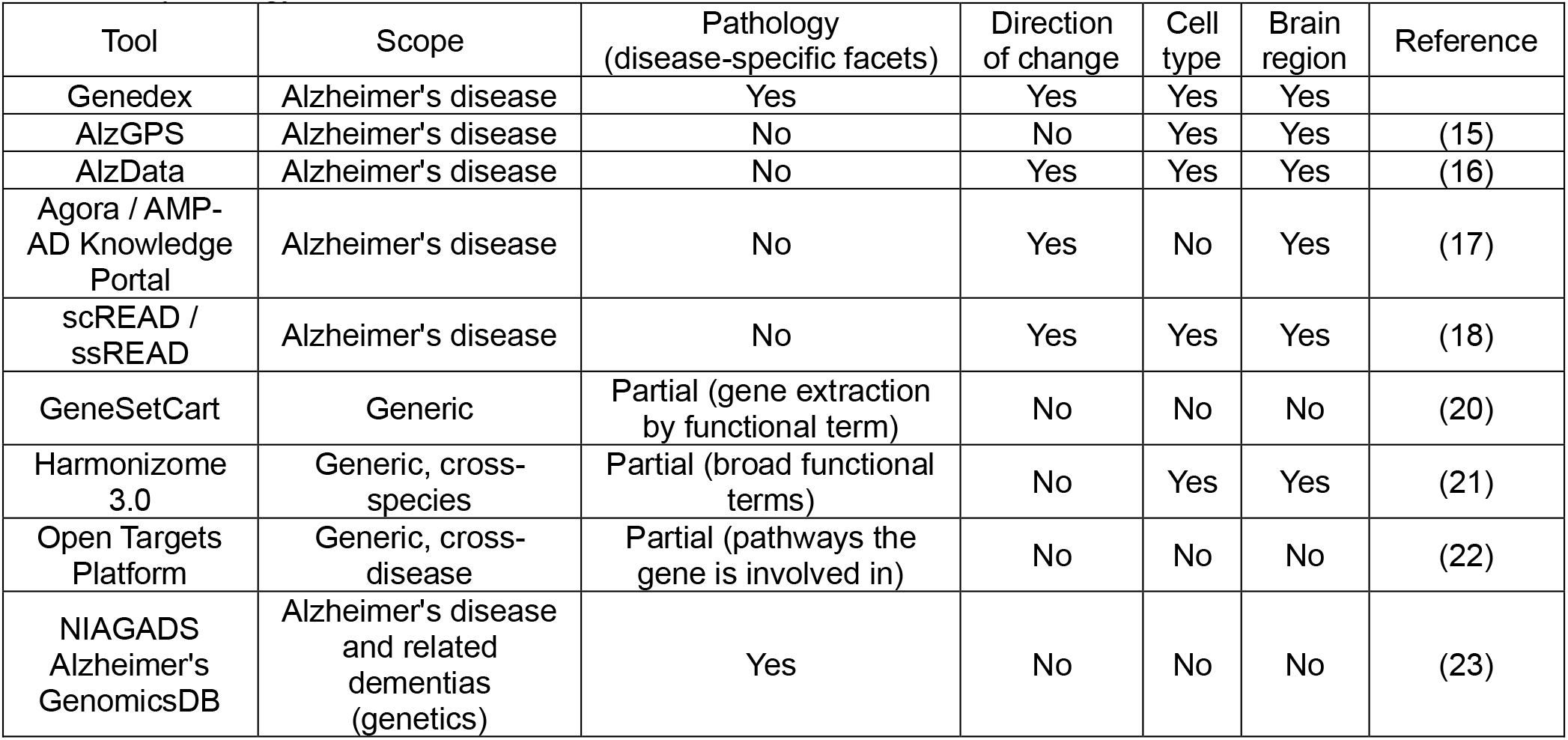
Coverage of the pathology, cell type, tissue, and direction of change required to position a gene within AD pathology.

| Tool | Scope | Pathology<br>(disease-specific facets) | Direction<br>of change | Cell<br>type | Brain<br>region | Reference |
| --- | --- | --- | --- | --- | --- | --- |
| Genedex | Alzheimer's disease | Yes | Yes | Yes | Yes |  |
| AlzGPS | Alzheimer's disease | No | No | Yes | Yes | (15) |
| AlzData | Alzheimer's disease | No | Yes | Yes | Yes | (16) |
| Agora / AMP-AD Knowledge<br>Portal | Alzheimer's disease | No | Yes | No | Yes | (17) |
| scREAD /<br>ssREAD | Alzheimer's disease | No | Yes | Yes | Yes | (18) |
| GeneSetCart | Generic | Partial (gene extraction<br>by functional term) | No | No | No | (20) |
| Harmonizome<br>3.0 | Generic, cross-<br>species | Partial (broad functional<br>terms) | No | Yes | Yes | (21) |
| Open Targets<br>Platform | Generic, cross-<br>disease | Partial (pathways the<br>gene is involved in) | No | No | No | (22) |
| NIAGADS<br>Alzheimer's<br>GenomicsDB | Alzheimer's disease<br>and related<br>dementias<br>(genetics) | Yes | No | No | No | (23) |

Genedex was developed to close the existing gap and contextualize genes within AD-relevant features (“facets”) using published gene signatures. Genedex indexes 1,562 curated signatures from 34 studies, each traceable back to its source publication. Each signature is annotated with facets representing different aspects of AD, such as the pathological process (e.g., “tau load” or “amyloid-β clearance”), cell type, tissue, and direction of change. Thus, each gene is represented by a profile across all four dimensions. Genedex is freely available as a web application (hidelab.github.io/genedex-app) built to interrogate any candidate gene along these dimensions.

We demonstrate the utility of Genedex through three studies, the first two of which test hypotheses arising from a recent study of cognitive resilience (6). First, we examine the expression of heat-shock chaperone genes, which have been proposed to enhance tau degradation, across tau-and amyloid-associated facets, to determine how their association with pathology varies by type of pathology, brain region, and cell type. Second, we profile *LINGO1*, a negative regulator of neurotrophic signaling in resilience, alongside *BDNF* and *NTRK2*, to examine how their relationship varies across pathological, regional, and cellular contexts. Third, we examine gene expression changes in vulnerable somatostatin-and parvalbumin-interneurons (19) to investigate how cellular vulnerability associates across the Genedex collection and whether it correlates with cognitive resilience. Genedex facilitates positioning genes within the broader context of AD facets. These investigations can generate novel hypotheses about disease mechanisms.

## Methods

### The Genedex Web Application

Genedex was implemented as a web app hosted on GitHub. hidelab.github.io/genedex-app. Users can query either an individual gene or a list of genes to examine their associations across functional, cellular, and tissue facets, together with the reported direction of change. For gene lists, pairwise similarity analysis and GenePrint (see below) visualization enable users to compare signatures and identify context-specific patterns across the curated collection.

### Facet taxonomy

Each gene list was annotated against a structured facet taxonomy. A facet describes one specific aspect of biological context, such as pathological/molecular process, cell type involved, tissue, or the genetic context. Facets are divided into four classes: functional, cellular, genetic, and tissue type. Functional facets describe molecular pathways and pathological processes, including amyloid-β production, aggregation and clearance, tau processing and load, neuroinflammation and microglial/astrocytic activation, synaptic and neuronal mechanisms, vascular and blood-brain barrier processes, and cognitive resilience and resistance (Supplementary Table 1). Cellular facets describe brain and peripheral cell types and states, spanning neuronal subtypes (excitatory and inhibitory neurons), astrocytes, oligodendrocytes, and oligodendrocyte precursor cells, microglial states (e.g., homeostatic, phagocytic, lipid-processing, inflammatory and disease-associated microglia); vascular cells (endothelial cells, pericytes, perivascular fibroblasts) and peripheral immune populations (Supplementary Table 2). Tissue-type facets record the anatomical source of the signature (e.g., prefrontal cortex, hippocampus, entorhinal cortex, blood) (Supplementary Table 3), and genetic facets (risk-increasing or protective) are defined for genomic variant-derived lists (Supplementary Table 4).

Facet definitions were mapped where possible to biomedical controlled vocabularies and ontologies, including Medical Subject Headings (MeSH), Mondo Disease Ontology (MONDO), Human Phenotype Ontology (HPO), and Gene Ontology (GO). Otherwise, facets were derived from primary literature. Every curated signature was required to map to at least one functional, one cellular, and one tissue facet.

Finally, each gene list was annotated with the direction of expression change wherever the source study reported one. Where a study reported both up-and downregulated genes, the two sets were stored as separate lists.

### Large language model-assisted curation

Signature discovery and ingestion were performed in a two-stage process supported by a large language model (LLM) configured with two fixed prompts. The first prompt, Genedex_Curator, took a single functional facet from the taxonomy as input, optionally restricted by cell type or tissue, and searched against PubMed, Rummagene, Gemma, and Dimensions (24,25) for candidate signatures. A report was generated that details the location of the gene list in the source (table, figure or supplementary material) and includes a score that ranked each candidate paper based on relevance to the input facet. A human curator then reviewed the report and downloaded the data for the selected candidates. The second prompt, Genedex_Ingestor, received the publication and the supplementary material and extracted the publication metadata (title, first author, journal, year, DOI, and PubMed identifier).

Each list was then assigned a sequential accession (GL followed by a seven-digit zero-padded number) and a descriptive name of the form GL(number)_(functional facet(s)_(cellular facet(s)_(directionality), where expression directionality was appended whenever applicable. All signatures were human, except murine disease-associated microglia signatures, which we included because of their importance to microglial biology.

Gene lists with more than 1,000 genes were truncated to the top 1,000 most significant genes.

### Gene list harmonization

Every existing curated gene list was checked against the current HGNC symbol database, producing a standardized copy of the entire collection using *HGNChelper* in R (v4.5.3). For each list, symbols flagged as HGNC-approved were retained unchanged, and symbols with a documented current replacement were renamed accordingly. Symbols with no current HGNC match were removed, and a filtering step separated unmatched symbols that resembled gene symbols from identifiers that are not gene name symbols, such as Ensembl and Havana clone identifiers and GenBank accessions.

### Gene search

*Gene search* takes a gene or a list of genes and finds every curated Genedex list in which they appear and returns gene lists from the Genedex collection that match the query gene(s). The user can filter the resulting gene lists for minimum overlap with the resolved query set and perform a statistical enrichment with a hypergeometric test. The resulting P-values were corrected for multiple testing across all lists tested using the Benjamini-Hochberg procedure. Results were returned as a detailed table, retaining functional and cellular facets as separate fields.

### Pairwise similarity analysis

Compositional similarity between gene sets was quantified using the Jaccard index, *J(A,B) = |A ∩ B| / |A ∪ B|*. Similarity was computed by gene list or associated facet. For gene lists, comparisons were made among selected user-defined and reference lists, optionally including search hits passing a user-defined adjusted p-value threshold. For AD facets, all lists sharing a given facet were first aggregated (union of their genes) to yield a facet-level set, and pairwise Jaccard indices were then computed among these aggregates. The resulting similarity matrices were rendered as heatmaps.

### GenePrint

*GenePrint* shows which AD processes, cell types, and brain regions each gene has been reported in, along with its direction of expression across a defined collection of studies. *GenePrint* can be constructed for each gene in the query. The Genedex collection can be restricted by direction of change, tissue of origin, study type and contrast, or limited to lists carrying a specified functional or cellular facet, permitting *GenePrint* to be computed within a defined biological subset; the displayed matrix may additionally be restricted to selected facet columns. Results are presented as genes x facets. The circle size scales with the number of lists in which the gene appears under each facet, and the color represents the directionality across the resulting lists. Blue represents consistently down-regulated genes, transitioning through grey for discordant directionality, to red for consistently up-regulated genes. The user can order rows by facet, by the order queried, or by hierarchical clustering, with each point highlighting the curated lists from which it derives.

### Application to resilience transcriptomic signatures

We assessed Genedex for resilience-associated genes extracted from Castanho & Naderi et al. (6). We first tested genes belonging to four heat shock chaperone families, HSP90/40/70/110, using GenePrint to investigate their association with tau-associated facets, which include tau load, tau aggregation, tau production, and tau degradation. Where functional facets overlapped with query heat shock genes, we stratified the results by brain region, showing the cell types associated with each functional facet. Regions were grouped to follow the anatomical sequence in which tau pathology spreads from the entorhinal cortex, through the hippocampus and inferior temporal gyrus, to association and primary cortical areas, so that successive panels correspond to successive stages of disease progression. We repeated the same analysis for amyloid-associated facets, grouping regions along the same axis so that the two sets of facets could be read against one another region by region. This grouping is applied to the amyloid facets for comparability with the tau panels. Exact panel membership differs slightly between facets because it was limited to the regions for which curated signatures were available or had a match. We then analyzed the functional association of *LINGO1*, testing it alongside *BDNF* and *NTRK2*.

### Pathway co-expression network

Although gene set relationships are conventionally assessed by gene overlap, such metrics fail to capture shared biological programs driven by distinct genes, a frequent occurrence when lists originate from different platforms, cell types, or statistical thresholds. Pathway Co-expression Network (PCxN)(26). PCxN summarizes the expression of each gene set in every sample of a dataset of human tissue transcriptomes. PCxN correlates the expression summaries between gene sets within each tissue and combines the tissue-level estimates into a single pathway correlation coefficient (PathCor) with an associated p-value (26). PathCor measures whether two gene sets co-vary in expression across samples. Two lists with no genes in common can be strongly correlated. PathCor ranges from −1 to 1. A positive coefficient indicates the two gene sets are expressed together. A negative coefficient indicates high expression of one set coincides with low expression of the other, as expected for programs active in different cell types or in opposing processes. Because the coefficients are estimated in normal tissue, they describe baseline co-regulation and do not report the direction of change in disease, which Genedex records separately for each list.

### Selection and clustering of correlated lists

Previously, Travaglini et al. (19) identified nine SST-positive and eight PVALB-positive interneurons that decline in abundance with AD progression. To examine their functional associations with the Genedex collection, we compared each signature against brain-derived Genedex lists. Lists outside the PCxN size range (5-1,000 genes) were filtered out. A threshold corresponding to the 99th percentile of absolute PathCor values across all PCxN gene-set pairs (excluding self-comparisons; |PathCor| = 0.757) was applied, resulting in 41 Genedex lists that crossed the threshold for at least one vulnerability list. All downstream analyses were performed in R (v4.5.3).

The resulting 41 x 34 PathCor matrix was clustered using Euclidean distance and Ward’s D2 linkage. Gene lists were clustered into 4 modules via dynamic tree cutting (minimum cluster size 5, deepSplit 2). Heatmaps were generated using *pheatmap* (v1.0.13), with rows annotated by Genedex functional and cellular facets, tissue, and direction of change, and columns annotated by interneuron subclass (SST or PVALB) and direction of change.

### Gene overlap

To verify gene overlap between the interneuronal vulnerability signatures and the 41 Genedex lists, we calculated the overlap coefficient, defined as the number of shared genes divided by the size of the smaller set. This metric was chosen to account for size disparities between the larger vulnerable signatures (median 853 genes) and the Genedex lists (median 228 genes). Overlap results were visualized using the row and column ordering of the PathCor heatmap.

### Shared pathway analysis

To characterize the shared functional processes of the vulnerability list-signature pairs, PCxN was rerun under identical parameters with the addition of the MSigDB 2024 collection (27). For each pair, MSigDB gene sets with an adjusted P < 0.05 and |PathCor| ≥ 0.50 with both members were included. Positively correlated pairs were ranked by their mean PathCor with both members, whereas anti-correlated pairs were ranked by mean absolute PathCor and classified by the sign of their correlation with each respective pair member.

## Results

### GFAP, TREM2 and APOE are recovered in their expected cell types and anticipated direction of change

To assess whether Genedex successfully recovers established AD biology, we first examined three well-characterized genes: *GFAP*, *TREM2*, and *APOE*. *GFAP* is upregulated in reactive astrocytes, which accumulate around amyloid plaques and increase with tangle burden (28). Genedex recovered the expected cell type and direction of change for all three genes (Supplementary Table 5).

*GFAP* has been widely reported to have higher gene expression in astrocytes in AD, including in single-nucleus transcriptomic studies (5). We therefore expected *GFAP* to map to astrocytic signatures and to be upregulated in AD. *GFAP* was detected in 61 gene lists, with astrocytes as the predominant cell type. It was upregulated in 16 of the 17 AD-associated astrocytic signatures that reported a direction of change, including seven amyloid plaque load signatures and one tau load signature.

*TREM2* encodes a microglial receptor and is required for a full microglial response to plaques (4,29). The rare *TREM2* R47H variant increases AD risk (30,31). *TREM2* was expected to map to microglial signatures and be upregulated with plaque load, microglial activation, and Aβ uptake. TREM2 was detected in 15 signatures, 13 of which came from microglia or macrophages. It was upregulated in all nine microglial differential expression signatures, which covered homeostatic, aging, antigen-presenting, amyloid-associated, and disease-associated (DAM) microglia. Four of these were Aβ-uptake signatures comparing immunized with non-immunized patients(32).

The ε4 allele of *APOE* is the strongest common genetic risk factor for late-onset AD (33). *APOE* is expressed by astrocytes but is induced in disease-associated microglia (4,34). In the entorhinal cortex, it has been reported to decrease in astrocytes and increase in a specific AD-associated microglial subpopulation (35). *APOE* was expected to map to microglial and astrocytic signatures, increasing in microglia but decreasing in astrocytes. *APOE* was upregulated in 39 of 40 microglial signatures, spanning AD, plaque load, tau load, and Aβ-uptake comparisons. In contrast, *APOE* expression was lower in resilient individuals than in AD in all three microglial and two of three astrocytic comparisons.

### Genedex combines curated signatures with a controlled vocabulary of associated AD facets

Genedex organizes AD gene signatures into a searchable resource that can be interrogated using a user-supplied gene list (**Figure 1A**). Each curated signature is stored as a separate gene list linked to the facets it is annotated with, so it can be found by name or through its facets. Genedex comprises 1,562 curated gene lists from 34 studies, annotated with four classes of facets: functional, cellular, brain region and genetic (**Figure 1B**). 28 functional facets describe molecular pathways and pathological processes such as amyloid-β production, aggregation and clearance, tau processing and load, neuroinflammation and glial activation, synaptic and neuronal mechanisms, vascular and blood-brain barrier processes, and cognitive resilience. 168 cellular facets describe neuronal, glial, vascular, and peripheral immune cell types and states. 25 tissue-type facets describe the anatomical source of each signature. Finally, genetic class catalogs variant-derived gene lists.

**Figure 1.**
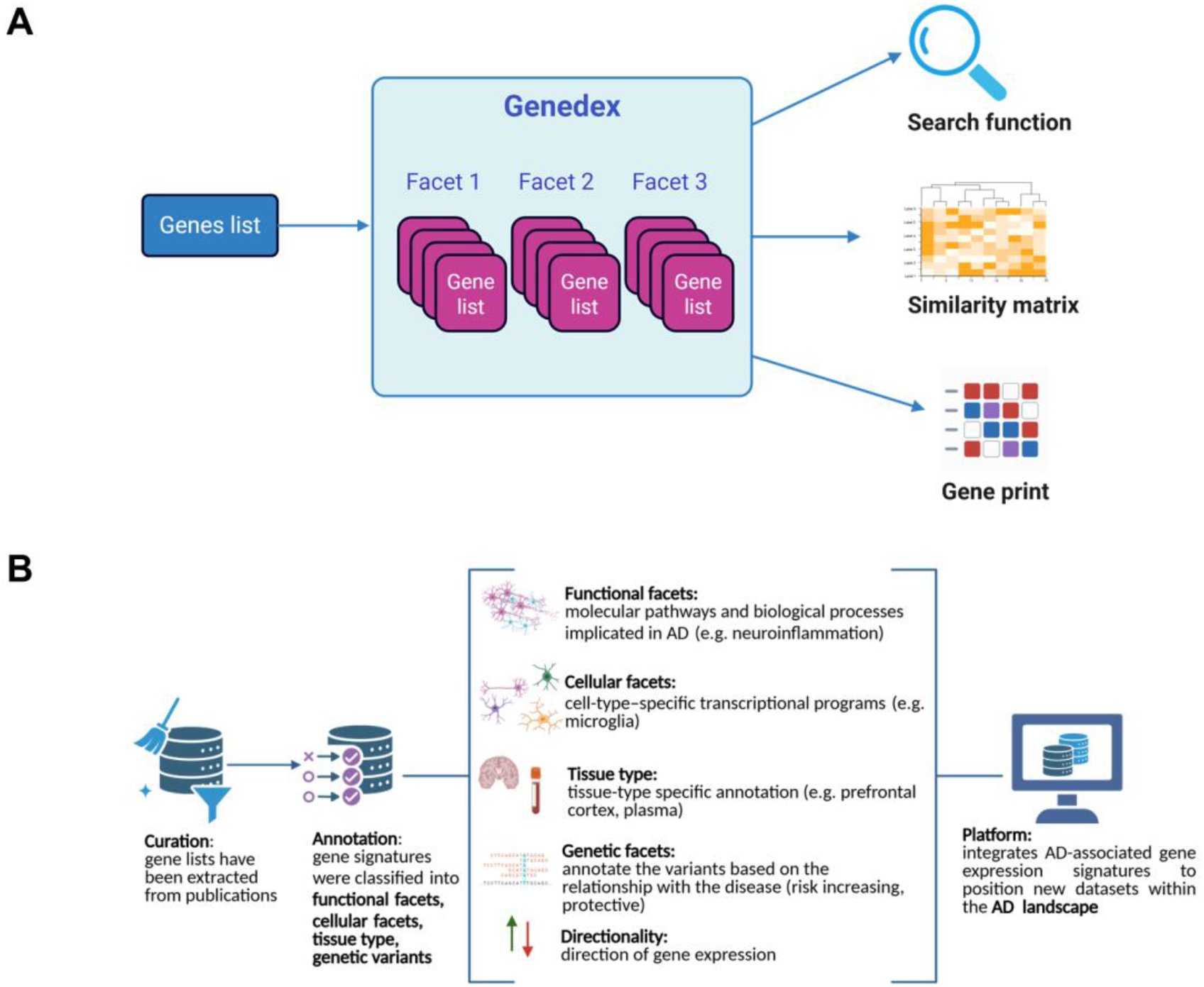
The Genedex framework. (**A**) Genedex is entered either through a facet or a user-supplied gene list. Three analytical functions operate on the organized collection: a search function, which searches for genes against the collection; a similarity matrix, which quantifies pairwise relationships among gene lists or facet-level; and a gene print, which tests which facets report a user-defined gene set more often than expected given the composition of the collection. (**B**) The curation and annotation pipeline. Gene lists are extracted from published studies, then annotated against four axes: functional facets (molecular pathways and pathological processes), cellular facets (cell-type-specific transcriptional programs), tissue type (anatomical source) and genetic facets (risk-or protective variants), together with the direction of change reported for each signature. The annotated collection is available through the platform, allowing genes to be positioned within the AD landscape.

Each curated signature has been mapped to at least one functional and/or cellular facet and one tissue-type facet, so every gene list has a defined biological context. The collection can be queried with three functions (**Figure 1A**). The search function matches any set of query genes, or a selected facet, against the curated collection and can test the overlaps for enrichment. A similarity matrix measures pairwise Jaccard similarity between gene lists, or between facets after pooling their lists, and orders them by hierarchical clustering to show which signatures and facets are most alike. GenePrint reveals which facets report a user-defined gene set more often than expected given the composition of the collection and shows both the strength of support and the reported direction of change across functional, cellular, and tissue-type facets.

Genes are frequently reported to change in opposite directions in different AD transcriptomic studies. These inconsistencies may reflect differences in biological context; for example, a gene can be upregulated in one cell type and downregulated in another or appear to change in opposite directions when studies make different comparisons. Because each list in Genedex has been annotated to its specific context, we tested whether facets account for these conflicting results.

Among the 1,272 lists from 19 studies that report a direction of change with disease and carry all three facet classes, 10,657 genes were upregulated in at least one list and downregulated in another (Supplementary Table 6). We added facet classes in sequence (functional, then cellular, then tissue) and considered a discordance explained when the two lists differed in the facet class added. A gene discordance was considered unexplained while any of its discordances occurred between lists that matched on all facet classes added so far. As a control analysis, we permuted facet labels among each study’s own lists. The proportion of genes with unexplained discordance was 72.9% after functional facets, 34.3% after cellular facets, and 12.9% after tissue facets. In permuted data, these proportions were 84.3% (95% range: 82.6-85.7%), 53.8% (95% range: 51.0-56.5%), and 29.5% (95% range: 26.0-33.1%).

These results show that discordant directionality in gene expression across independent studies does not necessarily reflect contradictions or evidence against a gene’s involvement in disease, but rather unexplained biological context. By generating a comprehensive facet taxonomy and annotating gene lists accordingly, Genedex provides a tool to contextualize directionality, resolving study conflicts and facilitating the interpretation of complex transcriptomic data.

### Heat-shock chaperones exhibit widespread upregulation with tau aggregation

To test our previous hypothesis that heat-shock chaperones enhance tau degradation (6), we evaluated the expression of 14 genes across four major heat-shock protein families (HSP70: *HSPA1A*, *HSPA1B*, *HSPA8*, *HSPA5*, *HSPA9*; HSP90: *HSP90AA1*, *HSP90AB1*, *HSP90B1*, *TRAP1*; HSP40: *DNAJA1*, *DNAJB1*, *DNAJB6*, *DNAJC5*; and HSP110: *HSPH1*) against four tau-associated pathology facets: aggregation, load, production, and degradation (**Figure 2a**, Supplementary Table 7). Tau production and degradation facets yielded no matches. Whereas tau aggregation and tau load showed widespread chaperone upregulation. For tau aggregation, nearly all chaperones were upregulated, except *HSPA8* and *TRAP1*, which yielded no matches, and *HSP90AB1*, which showed diverging directionality across cortical layers. Upregulation was similarly dominant for tau load, though a broader subset (*HSPA8*, *HSPA9*, *HSP90AA1*, *TRAP1*, *DNAJC5*, and *HSPH1*) displayed diverging directionality.

**Figure 2.**
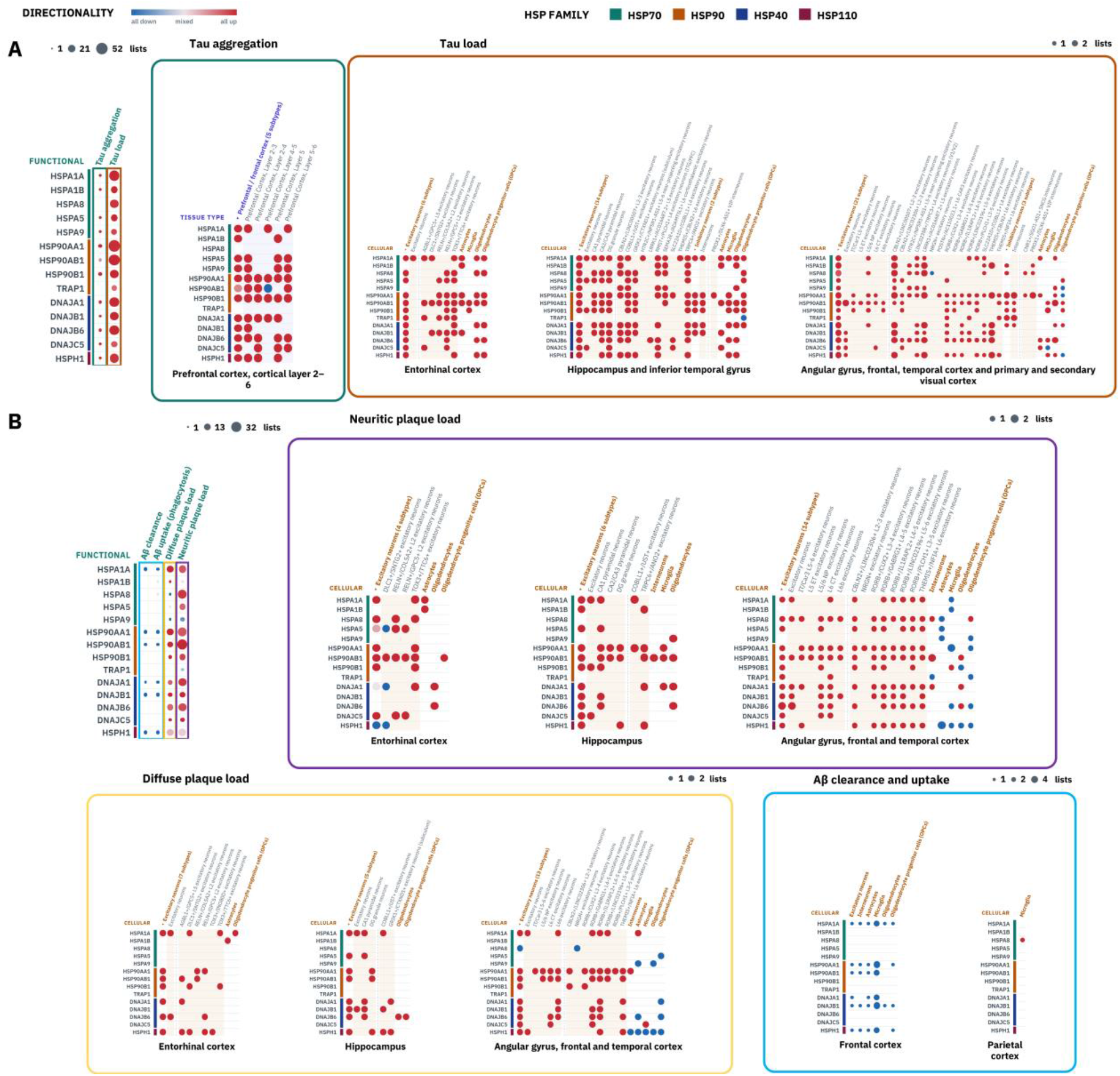
GenePrints of heat-shock chaperone genes across tau-and amyloid-associated facets. Fourteen genes from four heat-shock protein families are shown in every panel, grouped by the color bar on the left (HSP70: *HSPA1A*, *HSPA1B*, *HSPA8*, *HSPA5*, *HSPA9*; HSP90: *HSP90AA1*, *HSP90AB1*, *HSP90B1*, *TRAP1*; HSP40: *DNAJA1*, *DNAJB1*, *DNAJB6*, *DNAJC5*; HSP110: *HSPH1*). Dot size is the number of gene lists in which a gene appears under each facet, and color is its direction of change across those lists, from all down (blue) through mixed (grey) to all up (red). Facets boxed in the functional GenePrint on the left are resolved by tissue or cell type in the panel framed in the same color. (**A**) Tau-associated facets. Tau aggregation (green) is resolved by prefrontal cortical layer (layers 2-3 to 5-6), and tau load (orange) by cell type in three regional panels ordered by the sequence of tau spread: entorhinal cortex; hippocampus and inferior temporal gyrus; and angular gyrus, frontal, temporal, and primary and secondary visual cortex. (**B**) Amyloid-associated facets: Aβ clearance, Aβ uptake (phagocytosis), diffuse plaque load, and neuritic plaque load. Neuritic plaque load (purple) and diffuse plaque load (yellow) are resolved by cell type using the regional grouping in **A** (entorhinal cortex; hippocampus; and angular gyrus, frontal and temporal cortex), and Aβ clearance and uptake (blue) in frontal and parietal cortex.

Within the prefrontal cortex, chaperones differed in how many cortical layers showed upregulation (**Figure 2a**). *HSP90AA1* and *HSP90B1* were consistently upregulated across all cortical layers (layers 2-3 through 5-6). In contrast, other chaperones ranged from single layers (*HSPA1B* in layer 5; *DNAJB1* in layers 2-3) to four layers (*DNAJA1*, *DNAJB6*). *HSP90AB1* exhibited layer-specific discordance, being downregulated in layers 4-5 but upregulated in layers 2-4 and 5-6.

In regions affected early by tau pathology (the entorhinal cortex, hippocampus and inferior temporal gyrus), all 14 chaperones were upregulated with tau load (**Figure 2a**). The only exception was TRAP1 in hippocampal OPCs. The chaperone upregulation extended to regions affected later by tau pathology (angular gyrus, frontal, temporal and visual cortex). In those regions, *HSPA9*, *HSP90AA1*, and *HSPH1* were upregulated in OPCs; *DNAJC5* in microglia; and *HSPA8* in NRGN+ excitatory neurons.

### Amyloid pathology shows cell-specific transcriptional divergence and clearance-associated chaperone downregulation

Heat-shock chaperone genes were predominantly upregulated across brain regions and cell types with increasing tau burden. To evaluate whether the observed association was specific to tau pathology, we generated a GenePrint using amyloid facets. Aβ pathology revealed a more heterogeneous expression profile across diffuse plaque load, neuritic plaque load, Aβ clearance, and Aβ uptake (**Figure 2b**, Supplementary Table 8). For diffuse plaques, the angular gyrus, frontal, and temporal cortex panel exhibited a clear cell-type-dependent separation: *HSPA5*, *HSPA9*, *DNAJA1*, *DNAJB6*, and *HSPH1* were downregulated in glial and oligodendrocyte-lineage populations, whereas *HSPA1A*, *HSP90AA1*, *HSP90AB1*, *DNAJA1*, *DNAJB1*, *DNAJB6*, and *HSPH1* were upregulated in excitatory neurons. The hippocampus showed minimal involvement in this facet. Neuritic plaque load mirrored this glial-neuronal split in the angular gyrus, frontal, and temporal cortex panel; downregulation was concentrated in glial populations across ten chaperones, whereas seven chaperones were upregulated in layer-resolved excitatory neurons. The hippocampus did not show this separation; instead, three excitatory subtypes showed uniform upregulation (*HSPA5*, *DNAJA1*, and *HSPH1* in DLC1+/SNTG2+ excitatory neurons).

This cell-specific dichotomy shifted entirely when evaluating active Aβ handling rather than static plaque burden. During Aβ clearance and phagocytosis, the chaperone genes were predominantly downregulated. Chaperone genes such as *HSPA1A*, *HSP90AA1*, *HSP90AB1*, *DNAJA1*, *DNAJB1*, and *HSPH1*, which were upregulated alongside increasing plaque load in specific populations, were instead uniformly downregulated across nearly all frontal glial and neuronal populations, with the sole exception of *HSPA8* upregulation in parietal microglia. This widespread downregulation suggests that while chaperone genes are upregulated in response to amyloid burden, their expression is decreased during clearance processes. These results show that the chaperone response to amyloid pathology shows distinct cell-type-and region-specific transcriptional changes.

### LINGO1 shows convergent and divergent associations with the resilience markers BDNF and NTRK2 across cortex and midbrain

In our previous work (6), we identified *LINGO1* as a negative regulator of neurotrophic signaling in cognitive resilience. To understand its functional role in AD, we examined its expression profile alongside the canonical neurotrophic modulators *BDNF* and *NTRK2* using GenePrint (**Figure 3a**, Supplementary Table 9). We first characterized LINGO1 functionally in cortical regions, as we reported its downregulation in resilience cortical regions. *LINGO1*, *BDNF*, and *NTRK2* co-occurred in five facets: “Alzheimer’s disease”, “cognitive resilience”, “neuritic plaque load”, “selective vulnerability”, and “tau load”.

**Figure 3.**
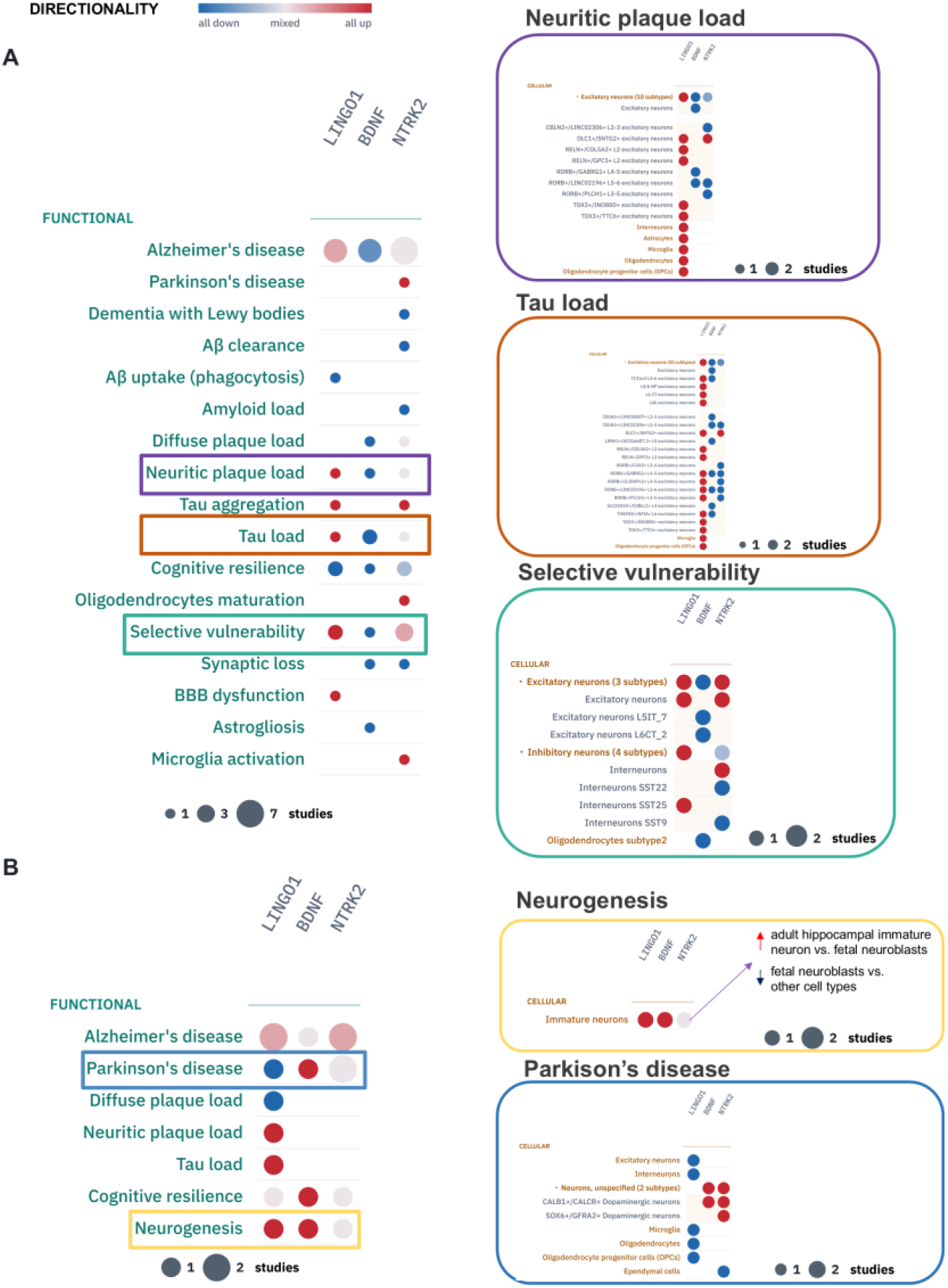
GenePrints of *LINGO1* and the resilience markers *BDNF* and *NTRK2* in cortex and midbrain. Circle size is the number of lists reporting a gene under each facet, and color is its direction of change across them, from all down (blue) through mixed (grey) to all up (red). Facets boxed in each functional GenePrint are resolved by cell type in the panel on the right, framed in the same color. (**A**) GenePrint with the collection restricted to cortical tissue signatures, with functional facets grouped into disease, amyloid pathology, and tau pathology. Neuritic plaque load (purple), tau load (orange), and selective vulnerability (green) are resolved by cell type, including layer-and marker-defined excitatory neuron subtypes and numbered SST interneuron subtypes. (**B**) GenePrint with the collection restricted to midbrain tissue signatures. Parkinson’s disease (blue) and neurogenesis (yellow) are resolved by cell type.

Single-cell resolution of pathology-associated facets revealed distinct cellular programs. Within the “neuritic plaque load” facet, *LINGO1* resulted as one of the most broadly expressed genes, showing uniform upregulation across all major glial populations (astrocytes, microglia, oligodendrocytes, and oligodendrocyte precursor cells), interneurons, and five of eight resolved excitatory neuron subtypes (*DLC1*/*SNTG2*, *RELN*/*COL5A2*, *RELN*/*GPC5*, *TOX3*/*INO80D*, and *TOX3*/*TTC6*). In the “selective vulnerability” facet, *LINGO1* was upregulated in excitatory neurons alongside *NTRK2*, whereas *BDNF* was downregulated in specific oligodendrocyte and excitatory populations (*L5IT_7*, *L6CT_2*). Among interneurons, *LINGO1* was selectively upregulated in the *SST25* subtype, whereas *NTRK2* showed bidirectional regulation across distinct somatostatin populations.

Against tau load, *LINGO1* co-occurred with *BDNF* and *NTRK2* in specific excitatory subtypes (*RORB*/*LINC02196* and *RORB*/*GABRG1*), with contrasting directionality: *LINGO1* was upregulated whereas *BDNF* and *NTRK2* were downregulated. Extending this analysis to midbrain tissue revealed a distinct, non-overlapping pattern of convergence across seven functional facets (**Figure 3b**, Supplementary Table 10). Within the “neurogenesis” facet, *LINGO1* and *BDNF* were co-upregulated in immature neurons (fetal hippocampal neuroblasts), reflecting an association of an early developmental state rather than active neurogenesis promotion. The three genes segregated by cell type in the “Parkinson’s disease” facet. *LINGO1* was broadly downregulated across glial and non-dopaminergic neuronal populations and notably absent from dopaminergic subtypes. Conversely, *NTRK2* and *BDNF* were specifically upregulated in dopaminergic neurons (*CALB1*/*CALCR* and *SOX6*/*GFRA2*), highlighting their role as dopaminergic markers rather than matching *LINGO1* expression patterns.

These results suggest that cortical *LINGO1* upregulation associates with pathological disease features, including selective vulnerability, tau load, and neuritic plaques, while contrasting with protective programs like cognitive resilience and Aβ uptake. In the midbrain, *LINGO1* aligns with immature developmental states and decouples from *BDNF* and *NTRK2*, demonstrating that its relationship with neurotrophic signaling varies by regional and cellular context.

### Separation of neuronal and glial programs by independent interneuron vulnerability

To evaluate whether cognitive resilience opposes neuronal vulnerability or operates within vulnerable populations, we examined inhibitory interneuron dynamics. Somatostatin (SST)-and parvalbumin (PVALB)-expressing interneurons exhibit selective loss in AD, whereas preservation of these neuronal populations has been associated with cognitive resilience (36–38). Using a multiregional single-nucleus dataset (19), we extracted SST and PVALB interneuron vulnerability signatures. Signatures from different studies, cell types, and statistical thresholds often share few genes even when they reflect related biology, so gene overlap alone can miss functional relationships between them. To capture these relationships, we sought to compare the interneuron vulnerability signatures with Genedex brain signatures at the pathway activity level, using Pathway Co-expression Network analysis (PCxN). Mapping 34 interneuron vulnerability signatures (comprising up-and downregulated expression programs across nine SST and eight PVALB supertypes) against 41 correlated Genedex signatures (|PathCor| > 0.757) revealed four distinct functional modules upon hierarchical clustering (Supplementary Figure 1 and Supplementary Table 11). Across the matrix, interneuron vulnerability signatures clustered by the direction of gene expression change. Genedex signatures clustered by cell type, revealing divergence between positively correlated neuronal programs and negatively correlated glial programs.

Clusters C2 and C3 represented positively correlated neuronal modules. Cluster C2 (n=9) displayed the strongest positive co-expression with vulnerable interneurons (**Figure 4a**), grouping selective vulnerability signatures across SST9 interneurons and upper/deep layer excitatory neurons alongside pathology-burdened neuronal states, hippocampal neural progenitors, and an entorhinal interneuron cognitive-resilience program (GL0000024). The correlation of resilience and vulnerability signatures within the same module suggests that transcriptional programs driving resilience may reflect cell-type-specific responses within vulnerable neuronal cells. Cluster C3 (n=8) formed another positively correlated neuronal module, dominated by pathology-load programs across specific excitatory subtypes (including TRPC6+/ANO2+, NRGN+ and CA2/CA3 populations) and entorhinal oligodendrocyte precursors.

**Figure 4.**
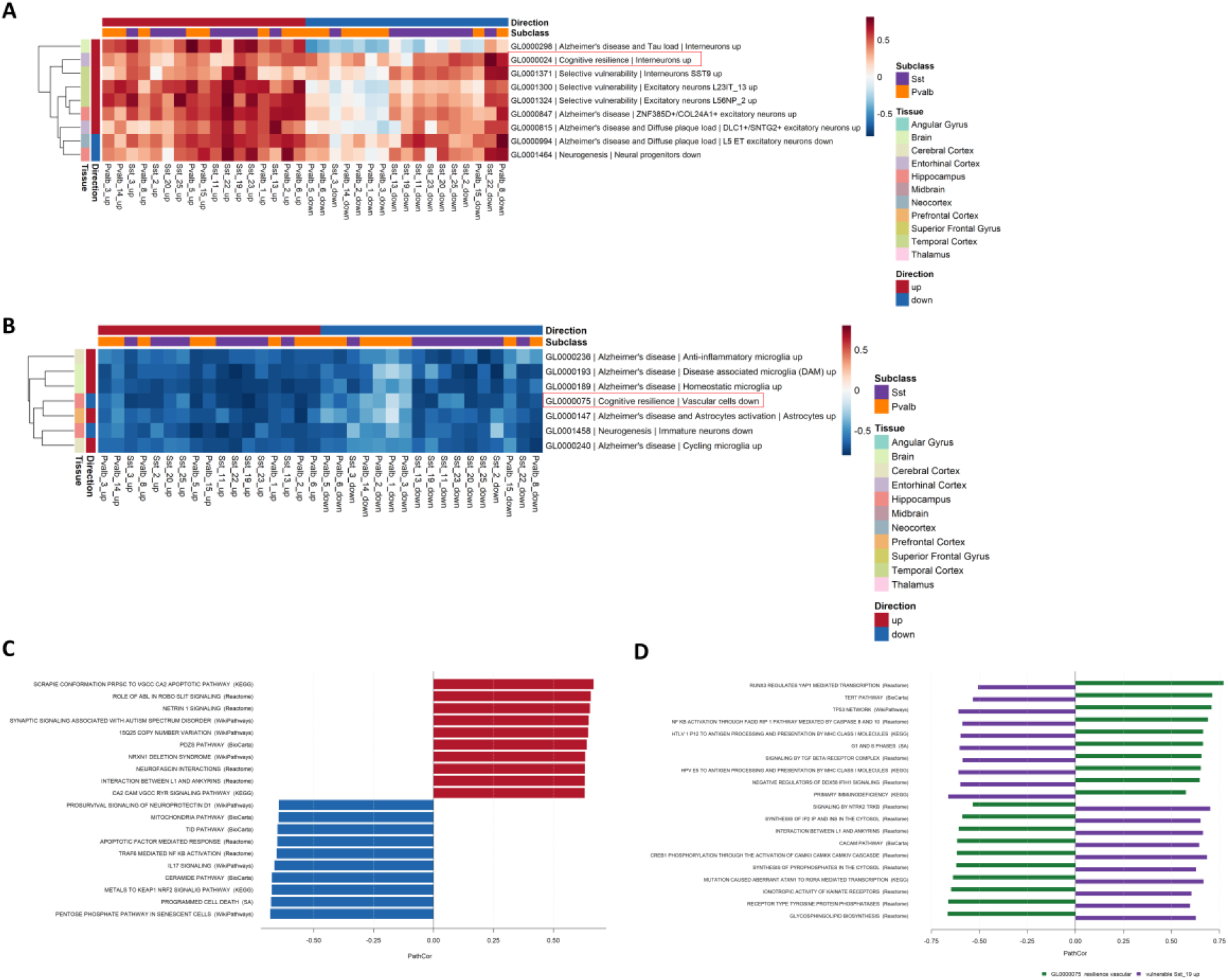
Pathway co-expression of Genedex signatures with vulnerable SST and PVALB interneurons. (**A, B**) Pathway correlation between Genedex brain signatures (rows) and the 34 interneuron vulnerability signatures from Travaglini et al. (columns): the up-and downregulated gene sets of nine SST and eight PVALB supertypes, annotated by direction of change and subclass. Rows are labeled by Genedex accession, functional facet, and cellular facet, and annotated by tissue and direction of change. Red boxes mark the two cognitive-resilience signatures: genes upregulated in interneurons (GL0000024) and genes downregulated in vascular cells (GL0000075) of resilient individuals compared with individuals with AD. (**C**) MSigDB pathways co-expressed with both members of the most positively correlated pair, GL0000024 and downregulated genes in SST22 vulnerable interneurons (PathCor = +0.780), showing the ten pathways with the highest and the ten with the lowest mean PathCor across the pair. (**D**) MSigDB pathways correlated with both members of the most negatively correlated pair, GL0000075, and upregulated genes in SST19 vulnerable interneurons (PathCor = −0.794). Bars show each pathway’s PathCor with GL0000075 (green) and SST19_up (purple) for the ten top-ranked pathways in each opposing pattern, ranked by mean

In contrast, Clusters C1 and C4 encompassed negatively correlated glial and non-neuronal programs. Cluster C1 (n=17) was the largest module, spanning AD-associated microglial, oligodendroglial, and astrocytic signatures alongside midbrain Parkinson’s disease programs. Cluster C4 (n=7) exhibited the strongest inverse correlations with vulnerable interneurons (**Figure 4b**), comprising diverse microglial functional states (homeostatic, disease-associated, anti-inflammatory, and cycling), activated astrocytes, immature neurons, and a hippocampal vascular cognitive-resilience signature, GL0000075, characterized by downregulated genes in resilient relative to AD brains. The uniform anti-correlation across distinct microglial states may indicate that this functional separation is dictated by cell type rather than specific glial activation phenotypes. Crucially, the two cognitive resilience programs mapped to opposing correlated clusters: entorhinal interneuron resilience correlated positively with neuronal vulnerability, whereas hippocampal vascular resilience correlated negatively.

Finally, a gene overlap analysis confirmed that the functional clusters were driven by pathway co-expression rather than direct gene overlap (Supplementary Figure 2 and Supplementary Table 12). Gene sharing across selected pairs was minimal, with the strongest positive pair sharing only 5 of 42 genes and the strongest negative pair sharing 81 of 1,000 genes.

### Interneuronal resilience genes correlate with synaptic pathways and against cell-death pathways

To identify the biological pathways underlying the strongest relationships identified by PCxN (**Figure 4b**), we determined which canonical MSigDB pathways were co-expressed with both members of the most strongly positively and negatively correlated signature pairs (**Figure 4c-d**). A positive correlation indicates that two gene sets are co-expressed, whereas a negative correlation indicates that high expression of one coincides with low expression of the other.

The most strongly positively correlated pair linked genes upregulated in entorhinal interneurons in resilient individuals compared with individuals with AD (GL0000024) with downregulated genes in the vulnerable SST22 supertype as AD pathology accumulates (PathCor = +0.780). Of 3,441 MSigDB pathways, 505 were correlated with both signatures (adjusted P < 0.05 and |PathCor| ≥ 0.50), and all of them correlated in the same direction with both signatures (Supplementary Table 13). The 102 pathways positively correlated with both signatures included synaptic and signaling pathways such as netrin-1 and ROBO SLIT signaling, PDZ scaffolding, neurofascin interactions, GABA-receptor signaling, and AMPA-receptor trafficking. Among the 403 negatively correlated pathways, programmed cell death, ceramide signaling, IL-17 signaling, TRAF6-mediated NF-κB activation, and intrinsic apoptosis were included (**Figure 4c**).

The most strongly negatively correlated pair linked downregulated genes in hippocampal vascular cells in resilient individuals compared with individuals with AD (GL0000075) with genes upregulated in the vulnerable SST19 supertype as AD pathology accumulates (PathCor = −0.794). Of the shared 132 pathways, 129 correlated in opposite directions with the two signatures (Supplementary Table 14). Of these, 97 were positively correlated with the vascular signature and negatively correlated with the SST19 signature, including TP53, G1 and S-phase, TGF-β-receptor, and NF-κB by FADD-caspase-8 signaling. The other 32 showed the reverse pattern, including NTRK2 TrkB signaling, CaMK-mediated CREB1 phosphorylation, L1-ankyrin interactions, and kainate receptor activity (**Figure 4d**). Only three pathways (interferon signaling, neutrophil degranulation, and GTSE1-mediated G2/M progression) were negatively correlated with both signatures. Pathway-level correlations show that synaptic and neurotrophic pathways associated with interneuron resilience are anti-correlated with the vascular genes reduced in resilience, whereas immune and cell-cycle pathways are positively correlated with these vascular genes. Evaluating interneuron vulnerability independently shows that cognitive resilience operates through cell-type-specific transcriptional networks that align with or oppose vulnerability, depending on cellular context.

## Discussion

Genedex integrates curated AD gene sets within a facet-structured vocabulary to enable systematic gene contextualization and hypothesis generation across distinct cell types and pathogenic mechanisms. A key advantage of Genedex is the annotation of transcriptional signatures by direction of change, along with cellular and regional context. By annotating transcriptomic data across these four classes, Genedex resolves discordant gene associations and supports the generation of hypotheses. Our findings on heat-shock chaperones, neurotrophic signaling regulators, and interneuron vulnerability demonstrate the utility of Genedex.

Our analyses reveal that chaperone gene regulation varies by pathology and cell type. Heat-shock chaperones were upregulated in tau-associated signatures across neurons and glia, broadly in entorhinal, hippocampal, and inferior temporal regions affected early by tau pathology (2), with an increase in protein-folding genes across multiple cell types in late-stage AD (39). Chaperone genes show cell-type-specific transcriptional differences in amyloid-associated facets. In plaque-load signatures, chaperones were upregulated in excitatory neurons and downregulated in glia, and in Aβ-clearance signatures they were downregulated almost uniformly. Because extracellular Hsp70 and Hsp90 promote microglial Aβ clearance (40), reduced glial expression may reflect diminished proteostatic support for Aβ handling. The exception was microglial *HSPA8*, which mediates clathrin uncoating and chaperone-mediated autophagy (41), a pathway that declines in AD (42).

Similarly, the *LINGO1* investigation highlighted the importance of annotating gene signatures by regional context. In the cortex, *LINGO1* was upregulated in selective-vulnerability, tau-load, and neuritic-plaque-load signatures and downregulated in cognitive-resilience and Aβ-uptake signatures. In tau-load signatures of *RORB*+ excitatory neuron subtypes, *LINGO1* was upregulated, whereas *BDNF* and *NTRK2* were downregulated, the inverse of the pattern we previously reported in cognitive resilience, in which *LINGO1* downregulation accompanied activation of BDNF-NTRK2 signaling (6). Given that LINGO1 promotes lysosomal degradation of activated Trk receptors (43), increased *LINGO1* and reduced *BDNF* and *NTRK2* expression may jointly limit neurotrophic signaling in tau-affected neurons. Studies in APP/PS1 mice support a detrimental role for LINGO1, where its antagonism improves cognition, enhances hippocampal neurogenesis and increases CB1R-rich GABAergic interneurons (44), promotes oligodendrocyte maturation (45) and reduces microglial activation and dendritic-spine loss (46), whereas hippocampal *LINGO1* overexpression impairs spatial memory and its silencing restores CB1R/TrkB signaling (47). Genedex also linked *LINGO1* to neurogenesis, but through its upregulation, together with *BDNF*, in fetal hippocampal neuroblasts and immature neurons. This may mark an immature neuronal state rather than a pro-neurogenic function, suggesting that LINGO1 acts differently during development and adult regeneration.

Mapping independent interneuron vulnerability against the Genedex collection delineated a functional separation between neuronal and vascular cells in relation to vulnerability. SST and PVALB signatures separated by direction of change rather than by subclass, which is consistent with vulnerability in AD being determined at the level of specific subtypes, including SST interneuron populations, and with the association between preserved inhibitory populations, including PVALB+ cells, and cognitive resilience (36–38). An entorhinal interneuron resilience signature clustered with neuronal programs positively correlated with interneuron vulnerability, in line with our previous observation that vulnerable SST populations are preserved in resilient individuals (6), and suggesting that resilience-associated expression can arise within vulnerable populations. This signature and the SST signature most strongly correlated with it were both positively correlated with synaptic and GABAergic pathways and negatively correlated with apoptotic and inflammatory pathways. As somatostatin regulates neprilysin-dependent Aβ degradation (48), preserved SST subtypes might contribute to amyloid processing.

Signatures anticorrelated with interneuron vulnerability were instead dominated by glial programs, suggesting that neuronal vulnerability and the glial response to pathology form distinct transcriptional compartments, consistent with the pathology-dependent remodeling of human microglia and astrocytes (5,49). A hippocampal vascular resilience signature, comprising genes downregulated in resilient relative to AD brains, was included within this compartment and was strongly anticorrelated with a vulnerable SST signature. Nearly all shared pathways correlated in opposite directions with the two, such as cell-cycle, TP53, TGF-β, and NF-κB-caspase-8 pathways aligned with the vascular signature, and neurotrophic and synaptic pathways, including TrkB signaling, aligned with the SST vulnerable signature.

### Limitations

Genedex is constrained by the literature from which it is curated, and therefore reflects existing biases in the regional, cellular, and disease-stage coverage of transcriptomic studies. Underrepresentation of a gene or facet should not be interpreted as evidence of absence. Facets also differ in curation depth, reflecting both curation effort and the extent to which biological processes have been studied. Additionally, multiple signatures may derive from the same study, cohort, or experiment. This is particularly relevant because approximately half of the gene lists currently in Genedex derive from a single study (50). This imbalance will diminish as further gene lists are added to the collection, and its effect on facet-level analyses is limited: only two functional facets (diffuse plaque load and neuritic plaque load) are represented exclusively by lists from this study. Furthermore, the underlying gene signatures are heterogeneous, having been generated using different experimental platforms, sample sizes, statistical thresholds, and biological contrasts. Finally, the analyses presented here are intended for contextualization and hypothesis generation, not causal inference.

### Future directions

Future work will extend coverage to further brain regions, cell types, disease stages, and other neurodegenerative conditions. Not every facet in the taxonomy is yet populated by at least one gene list, so continued curation will be needed to enrich the Genedex collection. We plan to integrate tools such as PCxN to identify pathway correlations from lists of interest for drug prioritization that emulate or inhibit PCxN-driven pathways.

## Data availability

The Genedex collection, metadata, and associated prompts are accessible at github.com/hidelab/Genedex. The Genedex web application is available for online use at hidelab.github.io/genedex-app, and the source code for local deployment can be downloaded from github.com/hidelab/genedex-app.

## Author Contributions

**Giulia Pegoraro**: Conceptualization, methodology, software, formal analysis, data curation, visualization, writing - original draft. **Mehmet Enis Isgoren**: Investigation, writing - review & editing. **Livia Rizzo**: Conceptualization. **Larisa Soto Morales**: Methodology, formal analysis. **Pourya Naderi Yeganeh**: Methodology, formal analysis, resources, writing - review & editing. **Inyoung Sung**: Methodology, formal analysis, writing - review & editing. **Sumeet Gupta**: Methodology, software, writing - review & editing. **Doo Yeon Kim**: Supervision, writing - review & editing. **Rudolph E. Tanzi**: Supervision, writing - review & editing. **Katie Lunnon**: Supervision, writing - review & editing. **Isabel Castanho**: Conceptualization, methodology, supervision, project management, writing - original draft, review & editing. **Winston Hide**: Conceptualization, methodology, supervision, project management, funding acquisition, writing - original draft, review & editing. Isabel Castanho and Winston Hide contributed equally to this work and share senior authorship.

## References

1. Thal DR, Rüb U, Orantes M, Braak H. Phases of Aβ-deposition in the human brain and its relevance for the development of AD. Neurology. 2002 Jun 25;58(12):1791–800. doi:10.1212/WNL.58.12.1791

2. Braak H, Braak E. Neuropathological stageing of Alzheimer-related changes. Acta Neuropathol (Berl). 1991 Sep;82(4):239–59. doi:10.1007/BF00308809

3. Arenaza-Urquijo EM, Vemuri P. Resistance vs resilience to Alzheimer disease: Clarifying terminology for preclinical studies. Neurology. 2018 Apr 10;90(15):695–703. doi:10.1212/WNL.0000000000005303

4. Keren-Shaul H, Spinrad A, Weiner A, Matcovitch-Natan O, Dvir-Szternfeld R, Ulland TK, et al. A Unique Microglia Type Associated with Restricting Development of Alzheimer’s Disease. Cell. 2017 Jun 15;169(7):1276–1290.e17. doi:10.1016/j.cell.2017.05.018

5. Habib N, McCabe C, Medina S, Varshavsky M, Kitsberg D, Dvir-Szternfeld R, et al. Disease-associated astrocytes in Alzheimer’s disease and aging. Nat Neurosci. 2020 Jun;23(6):701–6. doi:10.1038/s41593-020-0624-8

6. Castanho I, Naderi Yeganeh P, Boix CA, Morgan SL, Mathys H, Prokopenko D, et al. Molecular hallmarks of excitatory and inhibitory neuronal resilience to Alzheimer’s disease. Mol Neurodegener. 2025 Oct 1;20(1):103. doi:10.1186/s13024-025-00892-3

7. Jack Jr. CR, Andrews JS, Beach TG, Buracchio T, Dunn B, Graf A, et al. Revised criteria for diagnosis and staging of Alzheimer’s disease: Alzheimer’s Association Workgroup. Alzheimers Dement. 2024;20(8):5143–69. doi:10.1002/alz.13859

8. Abdulkhaliq AA, Kim B, Almoghrabi YM, Khan J, Ajoolabady A, Ren J, et al. Amyloid-β and Tau in Alzheimer’s disease: pathogenesis, mechanisms, and interplay. Cell Death Dis. 2026 Jan 9;17(1):21. doi:10.1038/s41419-025-08186-8

9. De Strooper B, Karran E. The Cellular Phase of Alzheimer’s Disease. Cell. 2016 Feb 11;164(4):603–15. doi:10.1016/j.cell.2015.12.056

10. Ashburner M, Ball CA, Blake JA, Botstein D, Butler H, Cherry JM, et al. Gene ontology: tool for the unification of biology. The Gene Ontology Consortium. Nat Genet. 2000 May;25(1):25–9. doi:10.1038/75556 PubMed PMID: 10802651; PubMed Central PMCID: PMC3037419.

11. Kramarz B, Roncaglia P, Meldal BHM, Huntley RP, Martin MJ, Orchard S, et al. Improving the Gene Ontology Resource to Facilitate More Informative Analysis and Interpretation of Alzheimer’s Disease Data. Genes. 2018 Nov 29;9(12):593. doi:10.3390/genes9120593

12. Wan YW, Al-Ouran R, Mangleburg CG, Perumal TM, Lee TV, Allison K, et al. Meta-Analysis of the Alzheimer’s Disease Human Brain Transcriptome and Functional Dissection in Mouse Models. Cell Rep. 2020 Jul 14;32(2):107908. doi:10.1016/j.celrep.2020.107908 PubMed PMID: 32668255; PubMed Central PMCID: PMC7428328.

13. Hou J, Hess JL, Zhang C, van Rooij JGJ, Hearn GC, Fan CC, et al. Meta-Analysis of Transcriptomic Studies of Blood and Six Brain Regions Identifies a Consensus of 15 Cross-Tissue Mechanisms in Alzheimer’s Disease and Suggests an Origin of Cross-Study Heterogeneity. Am J Med Genet B Neuropsychiatr Genet. 2025;198(4):e33019. doi:10.1002/ajmg.b.33019

14. Patel H, Dobson RJB, Newhouse SJ. A Meta-Analysis of Alzheimer’s Disease Brain Transcriptomic Data. J Alzheimer’s Dis. 2019 Apr 23;68(4):1635–56. doi:10.3233/JAD-181085

15. Zhou Y, Fang J, Bekris LM, Kim YH, Pieper AA, Leverenz JB, et al. AlzGPS: a genome-wide positioning systems platform to catalyze multi-omics for Alzheimer’s drug discovery. Alzheimers Res Ther. 2021 Jan 13;13(1):24. doi:10.1186/s13195-020-00760-w

16. Xu M, Zhang D, Luo R, Wu Y, Zhou H, Kong L, et al. A systematic integrated analysis of brain expression profiles reveals *YAP1* and other prioritized hub genes as important upstream regulators in Alzheimer’s disease. Alzheimers Dement. 2018 Feb;14(2):215–29. doi:10.1016/j.jalz.2017.08.012

17. Agora [Internet]. [cited 2026 Sep 7]. Available from: https://agora.adknowledgeportal.org

18. Wang C, Acosta D, McNutt M, Bian J, Ma A, Fu H, et al. A single-cell and spatial RNA-seq database for Alzheimer’s disease (ssREAD). Nat Commun. 2024 Jun 6;15(1):4710. doi:10.1038/s41467-024-49133-z

19. ravaglini KJ, Gabitto MI, Ding Y, Agrawal A, Postupna N, Mahoney JT, et al. Multiregional single-cell profiling reveals shared and specialized cellular vulnerability in Alzheimer’s disease. bioRxiv [Preprint]. 2026. doi:10.64898/2026.07.01.734821

20. Marino GB, Olaiya S, Evangelista JE, Clarke DJB, Ma’ayan A. GeneSetCart: assembling, augmenting, combining, visualizing, and analyzing gene sets. GigaScience. 2025;14:giaf025. doi:10.1093/gigascience/giaf025

21. Diamant I, Clarke DJB, Evangelista JE, Lingam N, Ma’ayan A. Harmonizome 3.0: integrated knowledge about genes and proteins from diverse multi-omics resources. Nucleic Acids Res. 2025 Jan 6;53(D1):D1016–28. doi:10.1093/nar/gkae1080

22. Buniello A, Suveges D, Cruz-Castillo C, Llinares MB, Cornu H, Lopez I, et al. Open Targets Platform: facilitating therapeutic hypotheses building in drug discovery. Nucleic Acids Res. 2025 Jan 6;53(D1):D1467–75. doi:10.1093/nar/gkae1128

23. Kuzma A, Valladares O, Greenfest-Allen E, Nicaretta H, Kirsch M, Ren Y, et al. NIAGADS: A data repository for Alzheimer’s disease and related dementia genomics. Alzheimers Dement. 2025;21(6):e70255. doi:10.1002/alz.70255

24. Zoubarev A, Hamer KM, Keshav KD, McCarthy EL, Santos JR, Van Rossum T, et al. Gemma: a resource for the reuse, sharing and meta-analysis of expression profiling data. Bioinformatics. 2012 Sep 1;28(17):2272–3. doi:10.1093/bioinformatics/bts430

25. Clarke DJB, Marino GB, Deng EZ, Xie Z, Evangelista JE, Ma’ayan A. Rummagene: massive mining of gene sets from supporting materials of biomedical research publications. Commun Biol. 2024 Apr 20;7(1):482. doi:10.1038/s42003-024-06177-7

26. Pita-Juárez Y, Altschuler G, Kariotis S, Wei W, Koler K, Green C, et al. The Pathway Coexpression Network: Revealing pathway relationships. PLoS Comput Biol. 2018 Mar 19;14(3):e1006042. doi:10.1371/journal.pcbi.1006042

27. Liberzon A, Subramanian A, Pinchback R, Thorvaldsdóttir H, Tamayo P, Mesirov JP. Molecular signatures database (MSigDB) 3.0. Bioinformatics. 2011 Jun 1;27(12):1739–40. doi:10.1093/bioinformatics/btr260

28. Serrano-Pozo A, Mielke ML, Gómez-Isla T, Betensky RA, Growdon JH, Frosch MP, et al. Reactive Glia not only Associates with Plaques but also Parallels Tangles in Alzheimer’s Disease. Am J Pathol. 2011 Sep 1;179(3):1373–84. doi:10.1016/j.ajpath.2011.05.047 PubMed PMID: 21777559.

29. Wang Y, Cella M, Mallinson K, Ulrich JD, Young KL, Robinette ML, et al. TREM2 Lipid Sensing Sustains the Microglial Response in an Alzheimer’s Disease Model. Cell. 2015 Mar 12;160(6):1061–71. doi:10.1016/j.cell.2015.01.049 PubMed PMID: 25728668.

30. Guerreiro R, Wojtas A, Bras J, Carrasquillo M, Rogaeva E, Majounie E, et al. TREM2 Variants in Alzheimer’s Disease. N Engl J Med. 2013 Jan 10;368(2):117–27. doi:10.1056/NEJMoa1211851

31. Jonsson T, Stefansson H, Steinberg S, Jonsdottir I, Jonsson PV, Snaedal J, et al. Variant of TREM2 Associated with the Risk of Alzheimer’s Disease. N Engl J Med. 2013 Jan 10;368(2):107–16. doi:10.1056/NEJMoa1211103

32. van Olst L, Simonton B, Edwards AJ, Forsyth AV, Boles J, Jamshidi P, et al. Microglial mechanisms drive amyloid-β clearance in immunized patients with Alzheimer’s disease. Nat Med. 2025 May;31(5):1604–16. doi:10.1038/s41591-025-03574-1

33. Corder EH, Saunders AM, Strittmatter WJ, Schmechel DE, Gaskell PC, Small GW, et al. Gene dose of apolipoprotein E type 4 allele and the risk of Alzheimer’s disease in late onset families. Science. 1993 Aug 13;261(5123):921–3. doi:10.1126/science.8346443 PubMed PMID: 8346443.

34. Krasemann S, Madore C, Cialic R, Baufeld C, Calcagno N, Fatimy RE, et al. The TREM2-APOE Pathway Drives the Transcriptional Phenotype of Dysfunctional Microglia in Neurodegenerative Diseases. Immunity. 2017 Sep 19;47(3):566–581.e9. doi:10.1016/j.immuni.2017.08.008 PubMed PMID: 28930663.

35. Grubman A, Chew G, Ouyang JF, Sun G, Choo XY, McLean C, et al. A single-cell atlas of entorhinal cortex from individuals with Alzheimer’s disease reveals cell-type-specific gene expression regulation. Nat Neurosci. 2019 Dec;22(12):2087–97. doi:10.1038/s41593-019-0539-4

36. Waller R, Mandeya M, Viney E, Simpson JE, Wharton SB. Histological characterization of interneurons in Alzheimer’s disease reveals a loss of somatostatin interneurons in the temporal cortex. Neuropathology. 2020;40(4):336–46. doi:10.1111/neup.12649

37. Mathys H, Peng Z, Boix CA, Victor MB, Leary N, Babu S, et al. Single-cell atlas reveals correlates of high cognitive function, dementia, and resilience to Alzheimer’s disease pathology. Cell. 2023 Sep;186(20):4365–4385.e27. doi:10.1016/j.cell.2023.08.039

38. O’Neill N, Stein TD, Olayinka OA, Empawi JA, Hu J, Tong T, et al. Cognitive resilience to Alzheimer’s disease characterized by cell-type abundance. Alzheimers Dement. 2024 Sep 11;20(10):6910–21. doi:10.1002/alz.14187 PubMed PMID: 39262221; PubMed Central PMCID: PMC12060125.

39. Mathys H, Davila-Velderrain J, Peng Z, Gao F, Mohammadi S, Young JZ, et al. Single-cell transcriptomic analysis of Alzheimer’s disease. Nature. 2019 Jun;570(7761):332–7. doi:10.1038/s41586-019-1195-2

40. Kakimura J, Kitamura Y, Takata K, Umeki M, Suzuki S, Shibagaki K, et al. Microglial activation and amyloid-beta clearance induced by exogenous heat-shock proteins. FASEB J. 2002 Apr;16(6):601–3. doi:10.1096/fj.01-0530fje

41. Bonam SR, Ruff M, Muller S. HSPA8/HSC70 in Immune Disorders: A Molecular Rheostat that Adjusts Chaperone-Mediated Autophagy Substrates. Cells. 2019 Aug 7;8(8):849. doi:10.3390/cells8080849

42. Bourdenx M, Martín-Segura A, Scrivo A, Rodriguez-Navarro JA, Kaushik S, Tasset I, et al. Chaperone-mediated autophagy prevents collapse of the neuronal metastable proteome. Cell. 2021 May 13;184(10):2696–2714.e25. doi:10.1016/j.cell.2021.03.048

43. Meabon JS, de Laat R, Ieguchi K, Serbzhinsky D, Hudson MP, Huber BR, et al. Intracellular LINGO-1 negatively regulates Trk neurotrophin receptor signaling. Mol Cell Neurosci. 2016 Jan 1;70:1–10. doi:10.1016/j.mcn.2015.11.002

44. He Q, Jiang L, Zhang Y, Yang H, Zhou CN, Xie YH, et al. Anti-LINGO-1 antibody ameliorates cognitive impairment, promotes adult hippocampal neurogenesis, and increases the abundance of CB1R-rich CCK-GABAergic interneurons in AD mice. Neurobiol Dis. 2021 Aug 1;156:105406. doi:10.1016/j.nbd.2021.105406

45. Yang H, Jiang L, Zhang Y, Liang X, Tang J, He Q, et al. Anti-LINGO-1 antibody treatment alleviates cognitive deficits and promotes maturation of oligodendrocytes in the hippocampus of APP/PS1 mice. J Comp Neurol. 2022;530(10):1606–21. doi:10.1002/cne.25299

46. Xie Y han, Jiang L, Zhang Y, Deng Y hui, Yang H, He Q, et al. Antagonizing LINGO-1 reduces activated microglia and alleviates dendritic spine loss in the hippocampus of APP/PS1 transgenic mice. Neurosci Lett. 2024 Jan 18;820:137612. doi:10.1016/j.neulet.2023.137612

47. He Q, Jiang L, Chao FL, Zhou CN, Zhang L, Zhou YN, et al. The role of LINGO-1 in regulating CB1R/TrkB signalling and GABAergic interneurons in Alzheimer’s disease pathogenesis. Exp Neurol. 2025 Sep 1;391:115319. doi:10.1016/j.expneurol.2025.115319

48. Saito T, Iwata N, Tsubuki S, Takaki Y, Takano J, Huang SM, et al. Somatostatin regulates brain amyloid β peptide Aβ42 through modulation of proteolytic degradation. Nat Med. 2005 Apr;11(4):434–9. doi:10.1038/nm1206

49. Smith AM, Davey K, Tsartsalis S, Khozoie C, Fancy N, Tang SS, et al. Diverse human astrocyte and microglial transcriptional responses to Alzheimer’s pathology. Acta Neuropathol (Berl). 2021 Nov 12;143(1):75–91. doi:10.1007/s00401-021-02372-6

50. Mathys H, Boix CA, Akay LA, Xia Z, Davila-Velderrain J, Ng AP, et al. Single-cell multiregion dissection of Alzheimer’s disease. Nature. 2024 Aug;632(8026):858–68. doi:10.1038/s41586-024-07606-7

